# GP2-enriched pancreatic progenitors give rise to functional beta cells *in vivo* and eliminate the risk of teratoma formation

**DOI:** 10.1101/2021.05.15.444293

**Authors:** Yasaman Aghazadeh, Farida Sarangi, Frankie Poon, Blessing Nkennor, Emily C. McGaugh, Sara S. Nunes, M. Cristina Nostro

**Author notes:** These authors contributed equally (Yasaman Aghazadeh, Farida Sarangi). Corresponding author: University Health Network, 101 College St. MaRS, PMCRT 3-916, Toronto, ON, M5G 1L7 (p).

## Abstract

Human pluripotent stem cell (hPSC)-derived pancreatic progenitors (PPs) can be differentiated into beta-like cells *in vitro* and *in vivo*, and therefore have therapeutic potential for type 1 diabetes (T1D) treatment. However, the purity of PPs varies across different hPSC lines, differentiation protocols and laboratories. The uncommitted cells may give rise to non-pancreatic endodermal, mesodermal, or ectodermal derivatives *in vivo,* hampering the safety of hPSC-derived PPs for clinical applications. Recently, proteomics and transcriptomics analyses identified glycoprotein 2 (GP2) as a PP-specific cell surface marker. The GP2-enriched PPs generate higher percentages of beta-like cells *in vitro* compared to unsorted and GP2^−^ fractions, but their potential *in vivo* remains to be elucidated. Here, we demonstrate that the GP2-enriched-PPs give rise to all pancreatic cells *in vivo*, including functional beta-like cells. Remarkably, GP2 enrichment eliminated the formation of teratoma *in vivo*. This study establishes that the GP2-enriched PPs represent a safe option for T1D treatment.

## Introduction

Type 1 diabetes (T1D) is an autoimmune disease characterized by the destruction of pancreatic beta cells. The standard treatment for T1D patients is the administration of exogenous insulin with frequent glucose monitoring, which significantly increases life expectancy. However, several risks associated with the disease remain prevalent; for example, patients undergoing insulin administration remain at high risk of hospitalization due to the development of long-term vascular complications such as stroke, retinopathy, nephropathy, neuropathy and cardiac disease (Aye and Atkin, 2014; Beck and Cogen, 2015). Therefore, beta cell replacement therapies that are able to endogenously regulate blood glucose are highly sought (Odorico et al., 2018; Shapiro et al., 2000; Shapiro et al., 2016). Indeed, islet transplantation using the Edmonton protocol has succeeded in controlling brittle diabetes, providing protection from hypoglycemic episodes and in several cases has led to insulin independence (McCall and Shapiro, 2012). In addition to donor-derived islets, the recent progress in the development of pancreatic cells from human pluripotent stem cells (hPSCs) offers the unprecedented opportunity to generate renewable, readily-available and quality-controlled pancreatic cells for clinical applications (Migliorini et al., 2021; Odorico et al., 2018). HPSCs can be differentiated into pancreatic progenitors (PPs) or pancreatic endodermal cells (PEC) which can give rise to functional beta-like cells both *in vitro* and *in vivo* (Kroon et al., 2008; Nostro et al., 2015; Rezania et al., 2014; Rezania et al., 2012). Remarkably, PECs are currently being tested in clinical trials for diabetes treatment (NCT02239354 and NCT03163511). However, preclinical studies reported the formation of teratoma in 15-50% of mice transplanted with hPSC-derived PEC/PP populations (Kelly et al., 2011; Kroon et al., 2008; Rezania et al., 2012). Therefore, designing strategies to purify the PP population is imperative to ensure the safety of this therapeutic approach.

The pancreatic secretory granule membrane major glycoprotein 2 (GP2) was previously identified as a specific cell surface marker of hPSC-derived PPs and which is co-expressed with PTF1A and NKX6-1 by the putative multi-potent pancreatic progenitor in the human developing pancreas (Ameri et al., 2017; Cogger et al., 2017). Purification and *in vitro* culture of GP2-enriched epithelial cells, obtained from human pancreas at 7 weeks of development confirmed that these cells give rise to acinar cells in which GP2 is upregulated, as well as ductal and endocrine cells in which GP2 is downregulated or silenced (Ramond et al., 2017). Similarly, *in vitro,* GP2-expressing hPSC-derived PPs down-regulate GP2 expression as they differentiate to beta-like cells (Ameri et al., 2017; Cogger et al., 2017). Collectively, these studies confirmed that the GP2-expressing PPs are committed to the pancreatic fate. However, there is a lack of *in vivo* data examining their commitment to the pancreatic fate. In this study, we explored the developmental potential of GP2-enriched PPs *in vivo* and demonstrated that GP2-sorted PPs give rise to endocrine and exocrine cells *in vivo* including beta cells with similar or improved functional efficacy compared to unsorted PP population. Furthermore, we demonstrated that GP2 sorting prevents teratoma formation *in vivo*. These findings indicate that GP2-enriched PPs represent a safe cell population with potential for clinical use.

## Results

### GP2 enrichment from hESC-derived PP

To assess the developmental potential of GP2-enriched PPs, H1 and H9 human embryonic stem cell (hESC) lines were differentiated *in vitro* to stage 4 PPs using previously published protocols (Korytnikov and Nostro, 2016) (Fig. 1A). At the end of stage 4, flow cytometric analysis was performed to assess the expression of NKX6-1, PDX1 and GP2 (Fig. 1B-G). Results indicated that both H1 and H9 cell lines generate more than 80% PDX1^+^/NKX6-1^+^ cells (Fig. 1B-D), while the H1 cells generated higher frequencies of GP2^+^ cells (~67%) compared to H9 cells (~31%) (Fig. 1E-G). These results demonstrated that the percentage of PDX1/NKX6-1 expressing cells may not correlate with that of GP2 (Fig. 1D, G). This is consistent with previous studies demonstrating the transient expression of GP2 in the multi-potent PPs and that not all NKX6-1^+^/PDX1^+^ cells express GP2 at the end of stage 4 (Cogger et al., 2017; Ramond et al., 2017). To assess the efficiency of PP purification, GP2 enrichment was performed at the end of stage 4 for both cell lines using magnetic-activated cell sorting (MACS) (Fig. 1H) (Cogger et al., 2017). The purity of the H1-derived GP2-enriched population (henceforth referred to as GP2^+^) was 88% (n=6), while the Flow-through (Ft) and the Not Sorted (NS) cells retained approximately 67% GP2-expressing cells (Fig. 1I, J). The quantification of GP2 mean fluorescent intensity (MFI) suggested significant enrichment of GP2^high^ expressing cells in the GP2^+^ population compared to Ft and NS fractions (Fig. 1K). Approximately 76% GP2^+^ cells (n=3) were obtained using H9-derived PPs, which was lower than the H1-derived GP2^+^ cells (Fig. 1L, M). Nevertheless, the MFI of the GP2^+^ cells was similar between the two lines (Fig. 1K, N). The percentage of GP2^−^ cells was significantly lower in the GP2^+^ fraction compared to NS and Ft in both H1 and H9-derived PPs (Supplemental Fig. 1A, B). Overall, these data suggest that the frequency of GP2^+^ cells and GP2 cell surface protein expression/cell were higher in the GP2^+^ fraction compared to NS and Ft populations (Fig. 1I-N). The recovery yield from MACS column was assessed by dividing the total number of GP2^+^ cells eluted from the MACS columns to the total cells loaded onto the columns and was 16% in both H1 and H9 lines (Table 1). Collectively, these data demonstrate the successful enrichment of GP2^+^ cells using MACS from both H1 and H9-derived PPs.

**Figure 1.**
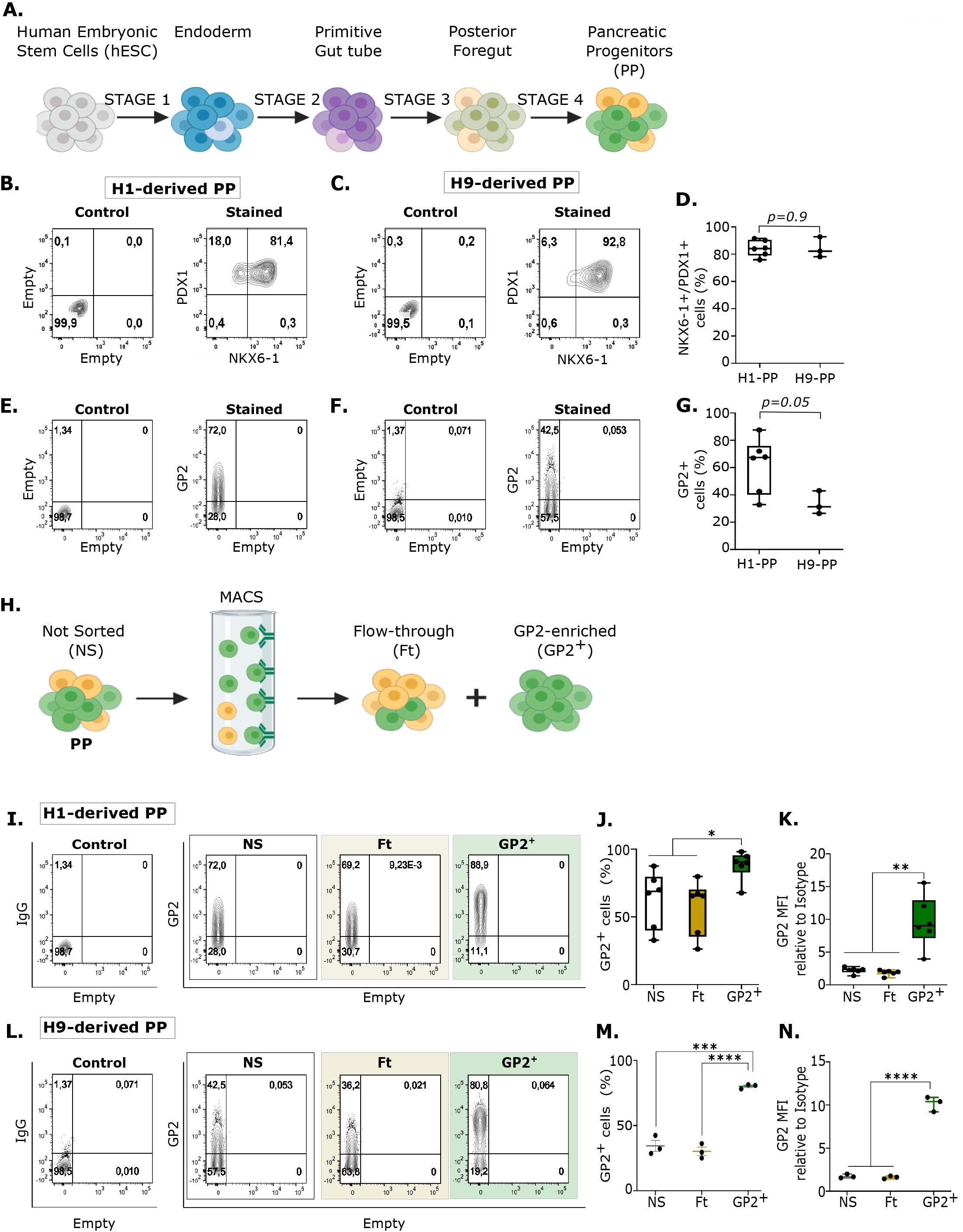
GP2 enrichment from hESC-derived PP. (A) Schematic of the hESC differentiation to pancreatic progenitors (PPs). (B-D) Representative flow cytometry plots and quantification of NKX6-1/PDX1 co-expression in the hESC-H1 (B) or H9 (C) derived PPs at the end of stage 4. (E-G) Representative flow cytometry plots and quantification of GP2 expression in H1- (E) or H9- (F) derived PPs at the end of stage 4 (G). (H1, n=6; H9, n=3; unpaired Student’s *T-test*; error bars represent S.E.M). (H) Schematic of GP2-enrichment using MACS. (I, J, L, M) Representative flow cytometry plots and quantification of the percentage of GP2 expressing cells in No Sort (NS), Flow-through (Ft) and GP2-enriched (GP2^+^) fractions of H1-(I, J) or H9-(L, M) derived PPs. (K, N) Mean fluorescent intensity (MFI) over isotype control in NS, Ft and GP2^+^ fractions of H1- (K) or H9- (N) derived PPs. (H1, n=6; H9, n=3; One-way ANOVA analysis with Tukey’s multiple test; *P<0.05, ** P< 0.01, *** P< 0.001, **** P<0.000; error bars represent S.E.M).

**Table 1.** GP2 recovery rate.

| Biological replicate | 1 | 2 | 3 | 4 | 5 | 6 |
| --- | --- | --- | --- | --- | --- | --- |
| % Cell recovery H1 | 27 | 10.86 | 11.81 | 6.65 | 10.4 | 28.16 |
| % Cell recovery H9 | 14.76 | 33.26 | 22.12 |  |  |  |

### Gene expression profile of GP2-sorted fractions

To better characterize the GP2^+^ cell population, we performed gene expression analysis of H1-derived GP2^+^, Ft and NS fractions by RT-QPCR. We confirmed that *GP2* expression was significantly higher in the GP2^+^ fraction compared to the Ft population, but not compared to the NS fraction, due to the high percentage of GP2-expressing cells present within the starting population (Fig. 2A). Further, we assessed the expression of PP markers *NKX6-1*, *PDX1* and *SOX9* and *PTF1A* (Schaffer et al., 2010; Zhou et al., 2007). No significant differences were detected in the levels of *NKX6-1*, *PDX1* and *SOX9* transcripts between GP2^+^, NS and Ft populations (Fig. 2B-E), while significantly higher *PTF1A* levels were detected in the GP2^+^ cells compared to the other populations (Fig. 2F). Since the expression of *PTF1A* is restricted to the acinar cells in the adult pancreas, we next assessed if the NS, Ft and GP2^+^ fractions show signs of early commitment to endocrine or exocrine lineages. We assessed the expression of acinar marker *CELA3* (Ramond et al., 2017; Schaffer et al., 2010) (Fig. 2F, G), ductal marker *HNF1B* (De Vas et al., 2015) (Fig. 2H) and endocrine markers *NEUROG3* and *NEUROD1* (Gu et al., 2002) in the three fractions (Fig. 2I, J), and detected no significant differences. Human cadaveric islets with ~80% purity were used as positive control. These data indicated that GP2-enriched cells express markers associated with PPs and sorting at stage 4 of differentiation does not enrich for exocrine or endocrine-committed progenitors. We further assessed if GP2 sorting could eliminate uncommitted cells from PPs by measuring the expression levels of neuroectodermal marker *ZIC3* (Bernal and Arranz, 2018) (Fig. 2K), mesodermal marker *PDGFRA* (Farahani and Xaymardan, 2015), (Fig. 2L), and pluripotency marker *OCT4* (Pan et al., 2002) (Fig. 2M) in the three fractions and compared them to hPSC-derived neuroectoderm, mesoderm, or undifferentiated hESCs respectively. These data indicated significantly lower marker expression in the NS, Ft and GP2^+^ fractions when compared to undifferentiated mesoderm, neuroectoderm or hESC controls, but no significant differences were observed between the three PP fractions (Fig. 2K-M). Since we achieved robust endodermal commitment (~99.2%) at stage 1 of H1 cells differentiation (Supplemental Fig. 2), the residual mesodermal, neuroectodermal or pluripotent cells might not be detected by RT-QPCR. Therefore, we next assessed the presence of non-pancreatic endodermal cells in the three fractions by assessing the expression levels of the anterior foregut markers *SOX2* and *NKX2-1* (Fig. 2N, O) and hindgut marker *CDX*2 (Fig. 2P). Although no significant differences were observed in *NKX2-1* expression between the three groups, the GP2^+^ sorted fraction demonstrated significantly lower *SOX2* and *CDX2* expression compared to NS and Ft fractions, respectively (Fig. 2N-P). These data demonstrate that GP2 is a reliable marker of multipotent PPs and that GP2 sorting may be used to eliminate non-pancreatic cells from PPs prior to transplantation.

**Figure 2.**
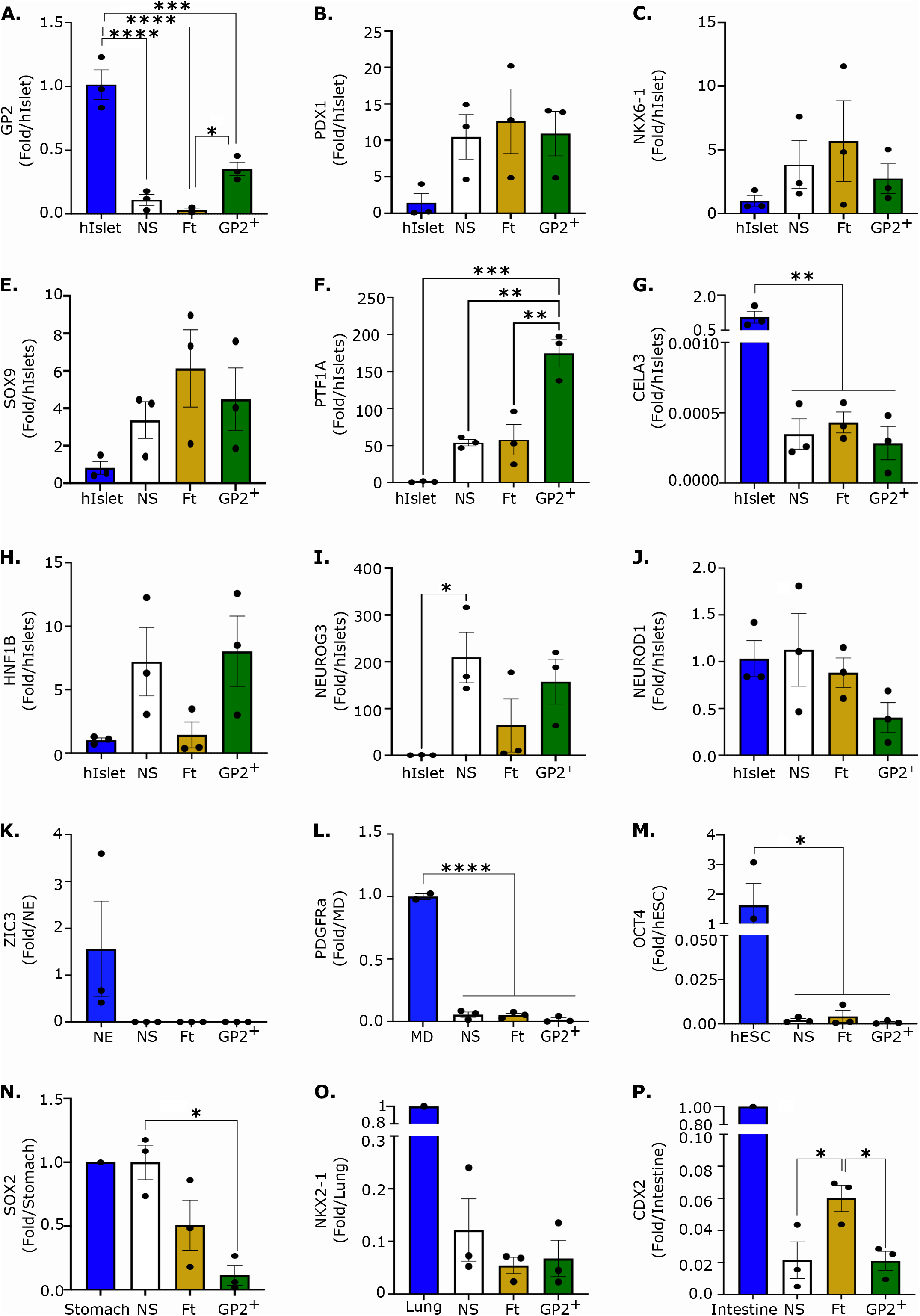
Gene expression profile of GP2-sorted fraction. RT-QPCR analysis for *GP2*, *PDX1*, *NKX6-1*, *SOX9*, *PTF1A*, *CELA3*, *HNF1B*, *NEUROG3*, *NEUROD1*, *ZIC3*, *PDGFRA, OCT4*, *SOX2*, *NKX2-1* and *CDX2* normalized to *RPL-19* housekeeping gene, in H1-derived NS, Ft and GP2^+^ PP fractions. hIslet is human islets; NE is neurectoderm; MD is mesoderm. (hIslets, NS, Ft, GP2^+^, NE, hESC, n=3; MD, n=2; human stomach, intestine and lung, n=1; One-way ANOVA and Tukey’s multiple test; *P< 0.05, ** P< 0.01, ***P<0.001, ****P<0.0001; error bars represent S.E.M).

### GP2 sorting prevents teratoma formation in vivo

We further assessed if GP2 sorting could reduce the risk of teratoma formation by transplanting the GP2^+^, NS and Ft cells in immunocompromised NOD.*Cg-Prkdc^scid^Il2rg^tm1Wjl^*/SzJ (NSG) mice. The cells were transplanted subcutaneously which allows teratoma detection with the naked eye. As the development of functional beta cells from transplanted PPs takes 12-18 weeks, we examined if GP2^+^, NS and Ft cells can survive in the subcutis long-term. We utilized an H1 line that constitutively expresses RFP (RFP-H1) (Davidson et al., 2012), differentiated them to PPs and further MACS sorted to generate GP2^+^, Ft and NS populations (Fig. 3A, Supplementary Fig. 3A). These populations were embedded in collagen hydrogels and transplanted subcutaneously into NSG mice. The RFP signal can be detected by live animal imaging, and therefore, we monitored the grafts *in vivo* at weeks 1, 4 and 12 post-transplantation (Fig. 3B). The background RFP signal was assessed using untransplanted mice (No Tx) and sham mice transplanted with empty hydrogels (Supplementary Fig. 3). RFP signal in NS, Ft and GP2^+^-transplanted mice was detected at all time points, confirming that all 3 populations can survive in the subcutis long-term. We utilized this model for the transplantation of MACS sorted H1-derived GP2^+^, Ft and NS populations, which were characterized by flow cytometry and QPCR (Fig. 1, 2). The transplanted NSG mice were monitored weekly and outgrowths visible with the naked eye were detected in NS and Ft recipients from week 8 onwards (Fig. 3C). Mice without visible outgrowths were maintained for 15 weeks, to allow for the development of functional beta cells (Kroon et al., 2008; Nostro et al., 2015; Rezania et al., 2014). The grafts were harvested and analyzed by Hematoxylin and Eosin (H&E) staining, which indicated the presence of mesodermal, ectodermal and endodermal cell types within the NS-(Fig. 3D) and Ft-(Supplementary Fig. 3A) derived grafts. Further pathological analysis using toluidine blue staining confirmed the presence of mesoderm-derived cartilage (Fig. 3E, Supplementary Fig. 4B), and immunostaining indicated the presence of SOX2 (stomach/neuronal marker, Fig. 3F, Supplementary Fig. 4C), FOXA2 (endodermal cells, Fig. 3G, Supplementary Fig. 4E), CDX2 (intestinal marker, Fig. 3H, Supplementary Fig. 4D), and NKX2-1 (lung marker, Fig. 3I, Supplementary Fig. 4F) expressing cells within NS and Ft-derived grafts. These data indicate that *in vivo*, the NS- and Ft-derived PPs give rise to non-pancreatic tissues from multiple germ layers, characteristic of teratoma (Kroon et al., 2008). Remarkably, none of the 11 mice transplanted with GP2^+^ cells developed teratomas/outgrowths while 27% (4/15) of NS recipients and 18% (3/17) of Ft-recipients developed teratomas/outgrowths (Fig 3J). These data strongly demonstrate that GP2 sorting prevents teratoma formation *in vivo.*

**Figure 3.**
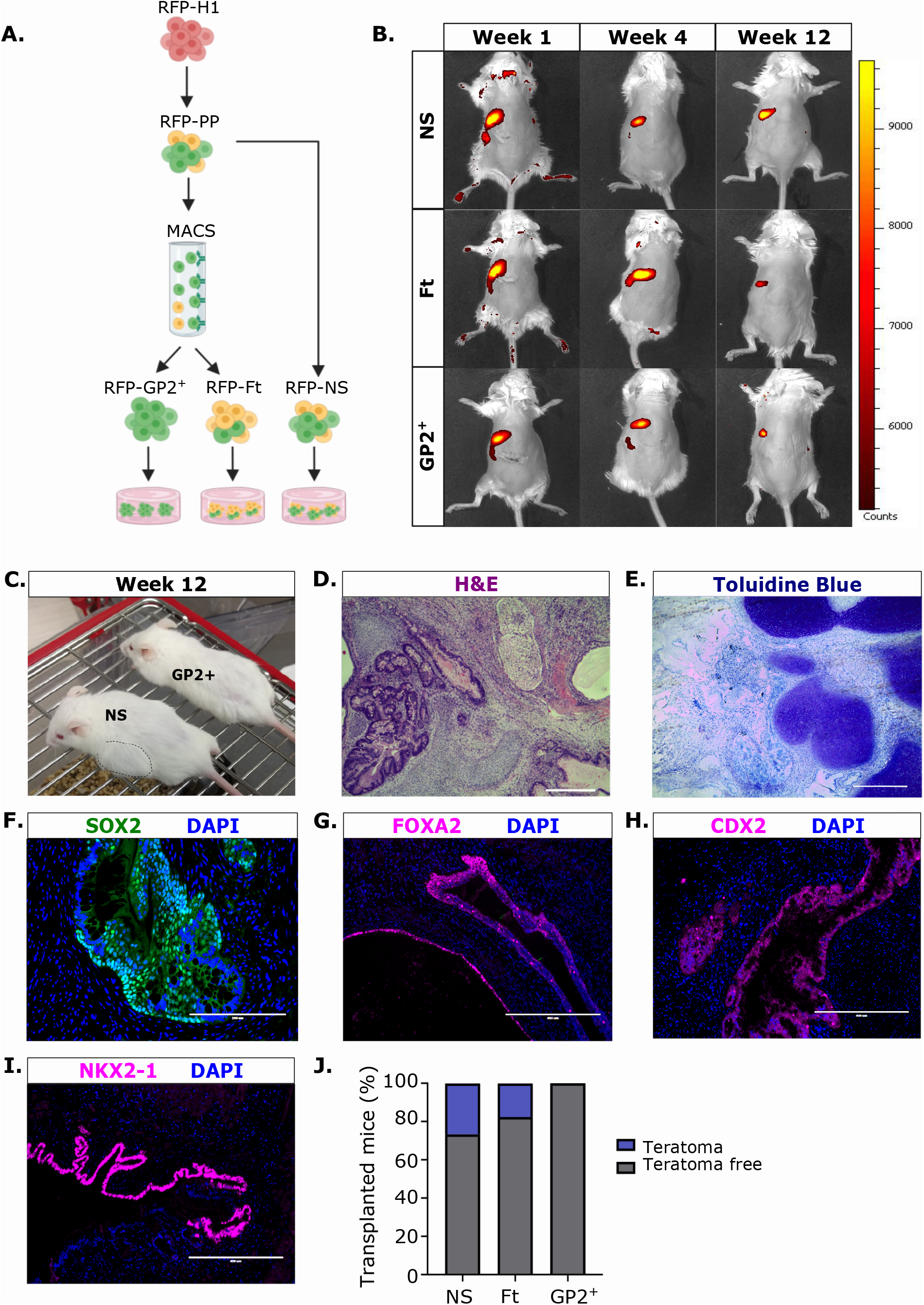
GP2 sorting prevents teratoma formation. (A) Schematic of RFP-H1 differentiation to PP followed by GP2-enrichment using MACS. (B) Live animal imaging of the NS, Ft and GP2^+^ at weeks 1, 4 and 12 post-subcutaneous transplantation. (C) Representative image of outgrowths visible with the naked eye in the mouse transplanted with NS cells (dashed line). No outgrowths detected in mice transplanted with GP2^+^ cells. (D-I) Representative images of NS-derived grafts stained with H&E (D), toluidine blue (E), or immunostained with SOX2 (F), FOXA2 (G), CDX2 (H) and NKX2-1 (F) antibodies at week 15 post-transplantation (D, scale bar is 100μm; F, scale bar is 200μm; E, G-I, scale bar is 400μm). (J) Percentage of NS, Ft and GP2^+^-recipient mice that generated teratomas/outgrowths (NS, n=15; Ft, n=17; GP2^+^, n=11).

### GP2^+^-enriched PPs give rise to smaller grafts compared to the unsorted fraction

Histological analysis of GP2^+^, NS and Ft grafts that did not generate teratomas at week 15 post-transplantation revealed graft size differences between the groups. This analysis indicated that significantly larger grafts are generated from NS population compared to the GP2^+^ population (Fig. 4A, B). To confirm, we evaluated the number of human cells within the graft by Ku80 immunostaining, a marker of human nuclei, which confirmed a significantly higher number of human cells in the NS grafts compared to the GP2^+^ grafts (Fig. 4C, D). To evaluate whether the NS fraction had a higher proliferative potential compared to the GP2-enrichd population, we quantified the expression of *MKI67* and *TOP2A* proliferation markers, in the MACS sorted populations by RT-QPCR and demonstrated significantly lower expression of both markers in the GP2^+^ fraction compared to NS fraction (Fig. 4E, F). The results confirmed that GP2^+^ cells are less proliferative than NS fraction, which could result in the formation of smaller grafts when transplanted *in vivo* (Fig. 4A-F).

**Figure 4.**
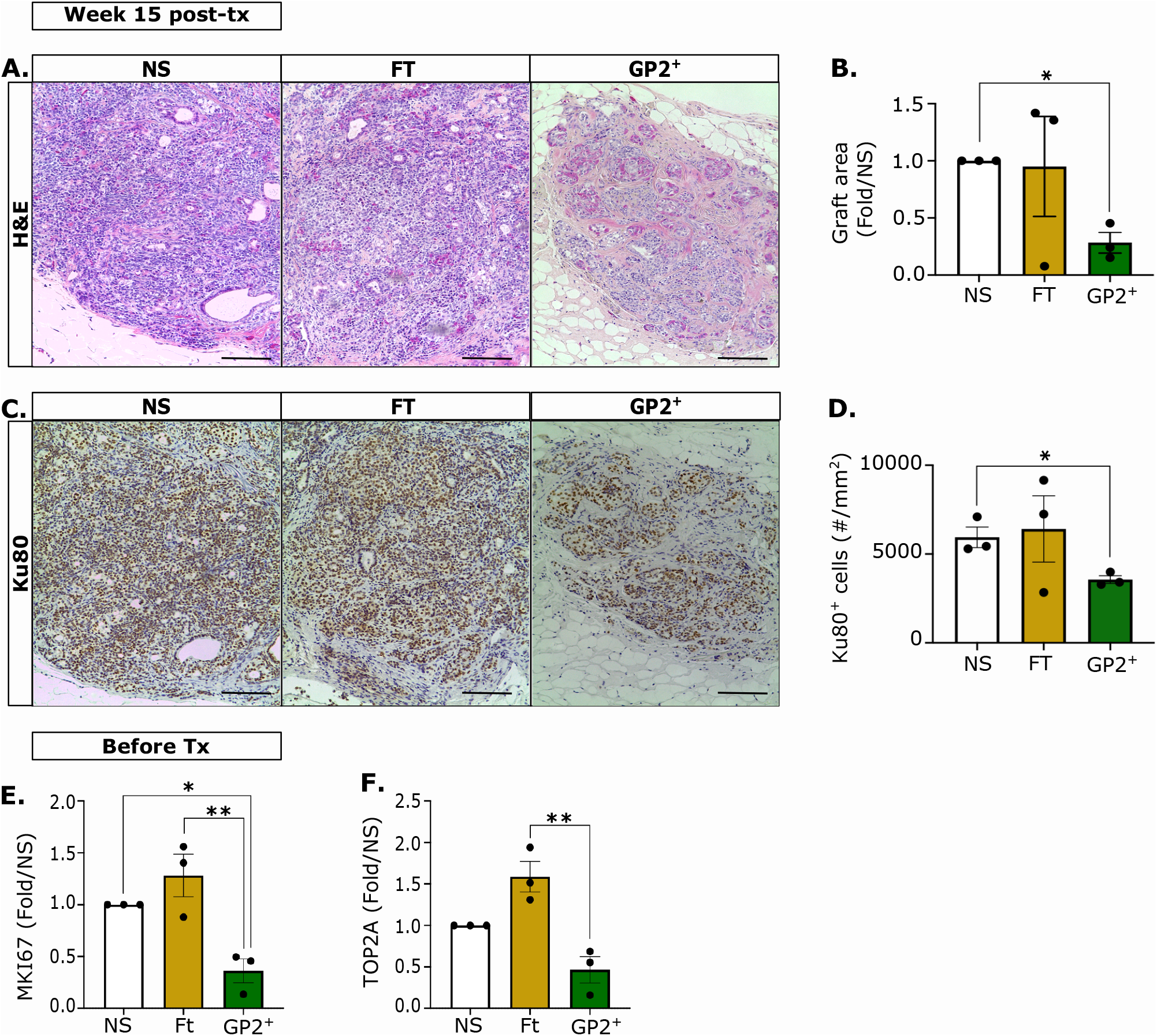
GP2-enriched PPs give rise to smaller grafts compared to the unsorted fraction. (A, C) Representative images of H&E or Ku80 staining of the non-teratoma grafts retrieved from mice 15 weeks post-transplantation (n=3, Scale bar is 400μm). (B, D) Quantification of graft area in panel A and Ku80^+^ nuclei in panel C. (mm^2^ is millimetre square; n=3; One-way ANOVA analysis with Tukey’s multiple test; *P< 0.05; error bars are S.E.M.). (E, F) RT-QPCR analysis of *MKI67* and *TOP2A* normalized to *RPL19* housekeeping gene in NS, Ft and GP2^+^ populations before transplantation. (n=3; One-way ANOVA analysis with Tukey’s multiple test; *P< 0.05, ** P<0.01; error bars are S.E.M.).

### GP2^+^-enriched PPs give rise to functional beta cells in vivo

We next examined whether GP2^+^ cells could differentiate *in vivo* to all the pancreatic lineages and retain the ability to generate functional beta cells. To assess functionality, glucose-stimulated C-peptide secretion assay was performed at week 15 post-transplantation, which indicated the presence of functional beta cells in GP2^+^, Ft and NS-derived grafts. Interestingly, the small size of the grafts derived from GP2^+^ cells did not jeopardize the concentration of secreted human C-peptide in response to glucose challenge (Fig. 5A). To assess reproducibility across stem cell lines, this experiment was replicated using H9-derived NS, Ft and GP2^+^ cells (Fig. 5B). Interestingly using this line, GP2^+^ recipients were the only group that secreted significantly higher C-peptide levels in response to glucose challenge which indicates that GP2 enrichment can be highly beneficial for PPs derived from hESC lines that have suboptimal GP2 expression at stage 4 (Fig. 1G). Immunofluorescence staining of the grafts at week 15 post-transplantation indicated that NS, Ft and GP2^+^ cells generated pancreatic endocrine cells (Chromogranin A, CgA), ductal cells (Cytokeratin 19, CK19) and acinar cells (Trypsin, TRYP) (Fig. 5C-H). Interestingly quantification of the number of acinar (TRYP^+^) cells indicated a significant reduction in the GP2^+^-derived grafts compared to the Ft-recipient grafts (Fig. 5E, G), while no differences in the total number of endocrine cells (CgA^+^) were detected (Fig. 5C, F). However, the ratio of endocrine to acinar cells was comparable between the conditions, suggesting that GP2 sorting does not affect lineage commitment (Fig. 5H). The endocrine cell population was further analyzed by immunofluorescent staining which identified the presence of beta cells (C-peptide^+^, Cp), alpha cells (Glucagon, GCG) and delta cells (Somatostatin, SST) in grafts derived from NS, Ft and GP2^+^ PP populations (Fig. 5I). Collectively, these data confirm that GP2^−^enriched cells retain the ability to give rise to all exocrine and endocrine pancreatic cells *in vivo,* including functional beta cells.

**Figure 5.**
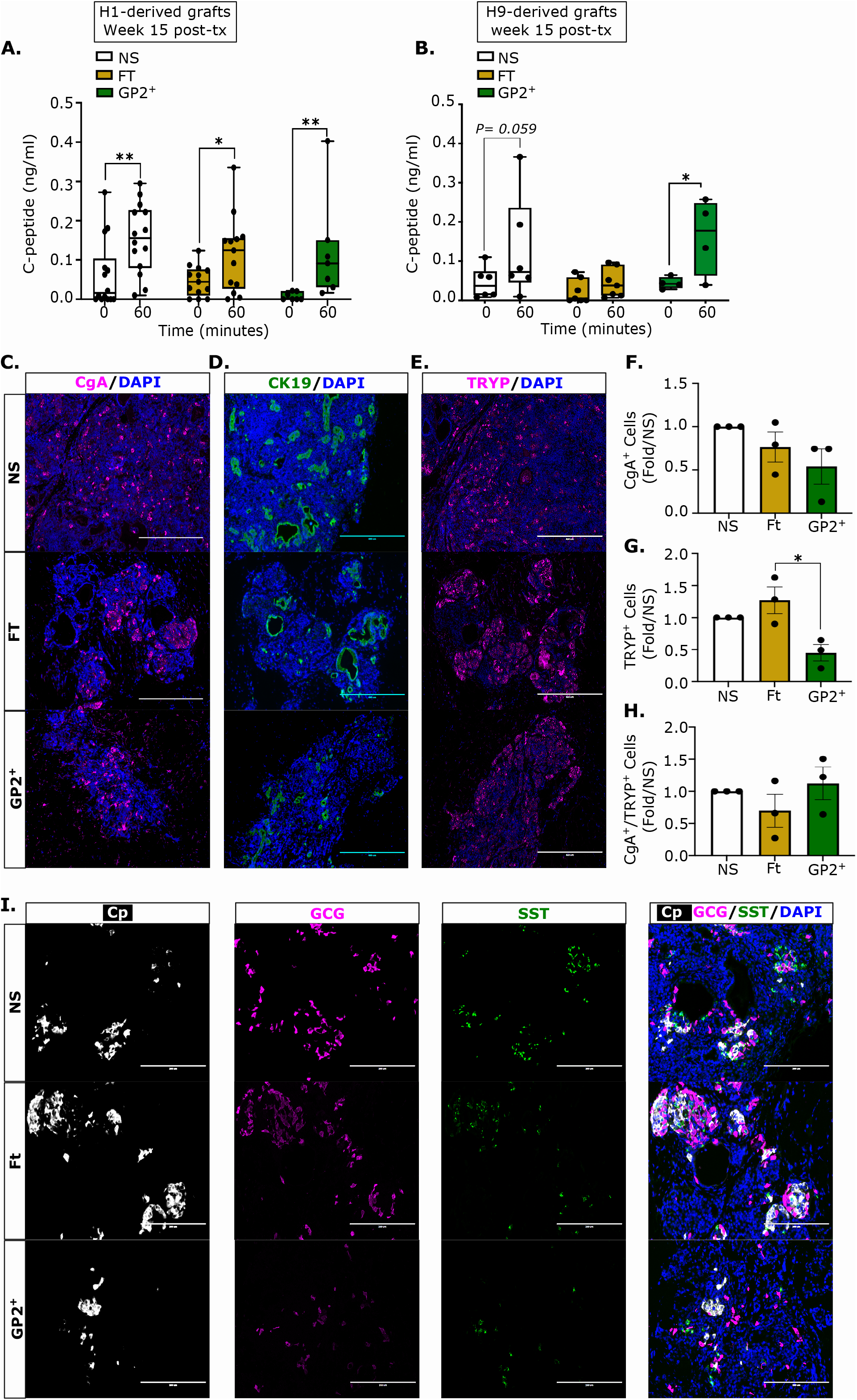
GP2^+^-enriched PP give rise to functional beta cells *in vivo*. (A, B) Glucose-stimulated C-peptide secretion assay at 15 weeks post-transplantation of H1- (A) or H9- (B) derived NS, Ft and GP2^+^ fractions (A: NS, n=15; Ft, n=14; GP2^+^, n=9; B: NS, n= 6; Ft, n= 7; GP2^+^, n= 4;Two-way ANOVA analysis with Tukey’s multiple test; *P< 0.05, ** P< 0.01; error bars indicate S.E.M.). (C-E, I) Representative immunostaining staining of grafts without teratoma retrieved from mice at week 15 post-transplantation using Chromogranin A (CgA, endocrine cell marker), Cytokeratin 19 (CK19, ductal cell marker), Trypsin (TRYP, acinar cell marker), C-peptide (Cp, beta cell marker), Glucagon (GCG, alpha cell marker), Somatostatin (SST, delta cell marker) antibodies. DAPI marks cell nuclei. (Scale bar is 200μm). (F-H) Quantification of the number of cells expressing CgA (F), Trypsin (G), ratio of CgA over Trypsin (H) normalized to NS population. (n=3; One-way ANOVA analysis with Tukey’s multiple test; *P<0.05; error bars are S.E.M.).

## Discussion

Stem cell-derived pancreatic cells are currently in clinical trials for beta cell replacement therapy with the goal to treat T1D. The hESC differentiation to PPs has variable yield between cell lines, protocols, and laboratories, and previous reports indicated the formation of teratomas from hESC-derived pancreatic cultures (Kelly et al., 2011; Kroon et al., 2008; Rezania et al., 2012), which can be an obstacle for clinical translation of these cells. To purify the hESC-derived PP population and eliminate uncommitted cells, we and others have identified PP cell surface markers such as GP2, CD142 and CD24 which offer the potential to purify the PP population (Ameri et al., 2017; Cogger et al., 2017; Jiang et al., 2011; Kelly et al., 2011; Ramond et al., 2017). However, both CD24 and CD142 are expressed in PDX1^+^ cells that lack NKX6-1 expression (Cogger et al., 2017; Jiang et al., 2011); in addition, CD24 is expressed in several lineages (Jiang et al., 2011) and CD142 is expressed by undifferentiated hESCs (Cogger et al., 2017), while GP2 is not expressed in hESC cells or polyhormonal cells that lack NKX6-1 expression (Cogger et al., 2017). In this study, we demonstrated that transplantation of GP2-enriched cells prevents teratoma formation in mice. Furthermore, the GP2-enriched PPs retain the developmental potential to give rise to exocrine and endocrine pancreas *in vivo* including functional beta cells.

Moreover, we demonstrate that sorting for GP2 can be effective in stem cell lines that give rise to GP2^low^ expressing cells at stage 4 such as H9 cell line used in this study. As GP2 expression is downregulated in the transition from multi-potent progenitor to the endocrine/ductal progenitor cells (Cogger et al., 2017; Ramond et al., 2017), we speculate that the majority of the H9-derived PDX1^+^NKX6-1^+^ GP2^−^ stage 4 cells have already transitioned to the endocrine/ductal progenitor. This finding is pursuant to previous reports, which indicated that GP2 enrichment can increase the efficacy of beta cell differentiation *in vitro* (Ameri et al., 2017; Cogger et al., 2017). Therefore, due to their developmental potential and safety, the GP2-enriched PPs show great promise for clinical use. However, it is important to note that GP2 enrichment using MACS resulted in significant cell loss (16% cell recovery), which may limit the use of MACS in large-scale manufacturing. This approach could be improved by developing better antibodies against GP2 and by optimizing the magnetic separation steps. Cell purification with other techniques such as FACS is more efficient, but also leads to the loss of target cells (Kelly et al., 2011), therefore it was suggested that improving the differentiation protocols could serve as an alternative to cell sorting. In this study, we show that even with robust differentiation of H1 line to endoderm at stage 1, and PPs at stage 4, the unsorted cells can give rise to teratomas, which could render the use of enrichment strategies imperative for clinical use. Nevertheless, this study shows that GP2 expression negatively correlates with the risk of tumorigenesis, and therefore the assessment of GP2 levels could be used for quality control prior to PP transplantation. Currently, NKX6-1 is prominently measured to assess pancreatic commitment at stage 4. This is based on mouse developmental studies which showed that NXK6-1 is the master regulator of beta cell fate (Sander et al., 2000; Schaffer et al., 2010; Taylor et al., 2013) and critical for the upregulation of insulin biosynthesis and glucose sensing genes (Aigha and Abdelalim, 2020; Schaffer et al., 2013). However, modeling pancreas development with hPSCs could generate aberrant cell types that co-express NKX6-1 and non-pancreatic transcription factors such as CDX2 and SOX2 (Veres et al., 2019; Wesolowska-Andersen et al., 2020). Such aberrant phenotypes are not unique to hPSC-derived pancreatic cultures and was reported in hPSC-derived lung and kidney populations (Hurley et al., 2020; Wu et al., 2018). Indeed, mapping hPSC-derived cell fate trajectories elucidated that early multipotent/progenitor cells could exhibit lineage plasticity (Petersen et al., 2017; Shih et al., 2015), suggesting that assessing the expression of a single cell marker does not accurately represent cell commitment. Furthermore, certain hPSC genetic backgrounds and differentiation protocols could exacerbate lineage plasticity of progenitor cells. In this study, we demonstrate that two hESC lines could give rise to similar levels of NKX6-1^+^/PDX1^+^ cells but vary significantly in the number of GP2^+^ cells. Therefore, in addition to NKX6-1, measuring GP2 expression could serve as an indicator of PP commitment and purity. Furthermore, recent studies identified additional cell surface markers such as CD177, expressed in endoderm at stage 1 (Mahaddalkar et al., 2020), and CD49a (ITGA1), expressed in beta-like cells at stage 6 that could provide further indication of successful differentiations (Veres et al., 2019). The CD177-enriched endoderm exhibited higher efficiency for pancreatic specification and differentiated to a homogenous NKX6-1^+^/PDX1^+^ at stage 4 *in vitro* (Mahaddalkar et al., 2020); however, whether transplantation of CD177^+^-derived PPs eliminates teratoma formation *in vivo* remains to be explored. Furthermore, to achieve an accelerated beta cell function *in vivo*, hPSC-derived beta cells can be generated *in vitro* and transplanted (Hogrebe et al., 2020; Pagliuca et al., 2014; Rezania et al., 2014; Velazco-Cruz et al., 2019). However, the yield of beta cell production is significantly lower than PP production (Cogger et al., 2017; Pagliuca et al., 2014; Rezania et al., 2014; Wesolowska-Andersen et al., 2020) which challenges scalability. Furthremore, the beta cells have a high oxygen consumption rate which could negatively impact survival post-transplantation (Papas et al., 2015; Pepper et al., 2012). Nevertheless, teratoma formation from hPSC-derived beta-like cells has not been reported, despite the presence of other endocrine and non-endocrine cells in hPSC-derived beta cell cultures (Veres et al., 2019). This could be attributed to the low proliferation rate of beta and non-beta cells present in these culture (McDonald et al., 2012; Oakie and Nostro, 2021; Rosado-Olivieri et al., 2020). Interestingly, in line with previous reports (Ameri et al., 2017), our data indicated that the GP2-enriched PPs are less proliferative than unsorted PPs and give rise to smaller grafts *in vivo* without jeopardizing graft efficacy in responding to glucose challenge. Additionally, similarly to the unsorted fraction, GP2-enriched PPs maintain the ability to give rise to all endocrine and exocrine cells. Therefore, GP2 enrichment generates a population with significantly lower non-pancreatic and proliferation marker expression which prevents teratoma formation *in vivo*, while maintaining potential for endocrine and exocrine pancreas development.

Noteworthy, previous studies indicated that the *in vivo* microenvironment can also influence cell fate specification and cell growth as the same cell population transplanted into different sites developed teratomas with various compositions and at different rates (Hentze et al., 2009; Lawrenz et al., 2004). Thus far, teratoma formation from unsorted PPs has been reported in the subrenal capsule (Kroon et al., 2008; Rezania et al., 2012) and epidydimal fat pad (Kelly et al., 2011; Kroon et al., 2008). In this study, we show teratoma formation in the subcutis, which is a clinically relevant site. However, it should be considered that encapsulation devices or other sites of transplantation expose the grafts to distinct microenvironments which could alter the risk of teratoma formation.

Collectively, our findings validate GP2 as a marker of committed PPs devoid of contaminants, which can give rise to functional beta-like cells *in vivo* and can be used to improve the safety of cell therapies.

## Experimental procedures

### Cell culture

H1 and H9 hESCs obtained from WiCell, were expanded *in vitro* to reach 70-80% confluency as previously described (Kennedy et al., 2007). The differentiation of hESC was carried out in 4 stages yielding definitive endoderm (D0-3), posterior foregut (D3-5), posterior foregut (D6-8) and pancreatic progenitors (D8-13).

### Magnetic Activated Cell Sorting (MACS)

At d13 of differentiation monolayers were dissociated to form single cells using TrypLE (Gibco) for 5 minutes at 37°C and washed with PBS without calcium and magnesium, 10% fetal bovine serum (FACS Buffer) containing DNase 1 bovine pancreas (10ul/ml, Millipore) and 10uM rock inhibitor Y27632 (Tocris). For cell surface GP2 staining, live cells were incubated with anti-GP2 antibody (1/10000 dilution, MBL) in FACS Buffer for 20 minutes at room temperature. Cells were washed with FACS buffer, followed by an incubation with anti-mouse PE-conjugated secondary antibody for 20 minutes at room temperature. Another wash with MACS buffer (Miltenyi Biotech) was performed using MACS buffer (75 ul /1×10^7^ cells) and PE magnetic beads were added (20ul/1 ×10^7^ cells). Cells and beads were incubated for 15 minutes at 4°C followed by a wash with MACS buffer. Cells were counted and NS fraction was used for further experiments. The remaining cells were loaded onto LS positive selection columns (Miltenyi Biotech), flow-through fraction was collected and treated with Y27632 prior to transplantation. The columns were washed twice with MACS buffer and cells were eluted in FACS buffer with Y27632. These cells constituted the GP2^+^ enriched fraction. These 3 fractions NS, FT and GP2^+^ were then analyzed for GP2 expression by flow cytometry and then cultured overnight at a plating density of 5×10^6^ cells/dish in Stage 5 media (+Y27632 and DNase1) to allow spontaneous aggregation before transplant. Based on the cell counts on day 13 of differentiation, 5 × 10^6^ cells were encapsulated in collagen hydrogels at day 14 for transplantation.

Neuroectoderm differentiation was adapted from previously described protocols (Chambers et al., 2009; Tchieu et al., 2017) and performed with the H1 cell line. Cells were cultured in Serum Free DMEM media (Gibco) and a mixture of DMEM (Gibco) with 15% KnockOut Serum Replacement (KSR; Gibco), 1mM Glutamine, and 1x non-essential amino acids (Gibco). Media changes were performed using the following base media: Day 0, 100% DMEM/KSR; Day 2, 100% DMEM/KSR; Day 4, 75% DMEM/KSR and 25% (Serum Free Differentiation Media) SFD (Korytnikov and Nostro, 2016); Day 6, 50% DMEM/KSR and 50% SFD; Day 8, 25% DMEM/KSR and 75% SFD; Day 10, 100% SFD. All media changes were supplemented with 10μM SB431542 (Sigma) and 500 ng/ml Noggin (R&D Systems) to promote neuroectoderm differentiation. All RNA samples were collected at Day 11. Neuronal differentiation was confirmed with immunofluorescence antibody staining for βIII-Tubulin (Chemicon, dilution: 1:1000,) and by RT-QPCR for neuronal markers (data not shown). Human islets were obtained from Alberta IsletCore (Lyon et al., 2016). Mesodermal cells were a gift from Dr. Stephanie Protze (McEwen Stem Cell Institute, University Health Network) and differentiated according to published protocol (Lee et al., 2017). Adult human stomach, intestine and lung mRNAs were purchased form Takara.

### Flow cytometry

At day 13 of differentiation monolayers were dissociated to form single cells using TrypLE (Gibco) for 5 minutes at 37°C. For intracellular NKX6.1 and PDX1 staining, cells were fixed in cytofix/cytoperm (BD Bioscience) for 24 hours at 4°C, washed twice with perm/wash (BD Bioscience) and resuspended in unconjugated PDX1 (1/100, R&D Systems) and NKX6.1 (1/1500, Developmental Studies Hybridoma Bank F55A10) antibodies overnight at 4°C. After washing two times with perm/wash the cells were incubated with a Donkey anti mouse Alexa-647 and a donkey anti-goat Alexa-488 (1/400, Jackson ImmunoResearch Laboratories) secondary antibodies for 20-30 minutes at room temperature. For cell surface GP2 staining, live cells were first incubated with anti-GP2 antibody (1/10,000) in 1× PBS with 10% fetal bovine serum (FACS Buffer) for 20 minutes at room temperature. Cells were washed once with FACS buffer and incubated for an additional 20 minutes at room temperature with a donkey anti-mouse Alexa-647-conjugated secondary antibody (1/400, Jackson ImmunoResearch Laboratories). After washing with perm/wash, flow cytometric analysis was performed on a BD LSR Fortessa flow cytometer and Flowjo V.10.1 was used for data analysis.

### Animal transplantation

After MACS sorting and aggregation overnight, 5 × 10^6^ GP2^+^ cells, NS cells or Ft cells were embedded in a collagen hydrogel composed of 3 mg/ml rat tail collagen type I, water, low glucose DMEM, Sodium bicarbonate and HEPES sodium salt. All animal procedures were performed according to the University Health Network guidelines. 4-6-week-old male NSG mice were anaesthetized using isoflurane. A subcutaneous incision was formed through which a collagen hydrogel was transplanted.

### Glucose-stimulated C-peptide secretion assay and ELISA assay

At week 15 post-transplantation, the animals were fasted overnight. The next day blood was collected in an EDTA coated tube though saphenous vein. Each mouse was weighed and 3g/kg glucose was injected intraperitoneally per mouse; 60 minutes later a second blood collection was performed from the saphenous vein. High-speed centrifugation was performed to separate the serum fraction. Ultra-sensitive C-peptide ELISA assay (Mercodia) was performed to measure the levels of human C-peptide in the sera.

### Live animal imaging

Mice were anaesthetized using isoflurane and live imaging was performed by PerkinElmer Xenogen IVIS Spectrum Imaging System.

### Immunostaining

Immunostaining was performed as previously described (Cogger et al., 2017). Briefly, grafts were embedded in agarose and paraffin, and 3 μm sections were cut by the Toronto General Hospital, Pathology Research Program Laboratory. Sections were de-paraffinized using xylene, and rehydrated in a serial dilution of absolute alcohol. Antigen-retrieval was performed, and the sections were blocked using 10% non-immune donkey serum (Jackson ImmunoResearch Laboratories Inc.) in PBS. Primary antibodies were diluted in 1X PBS supplemented with 0.3% Triton X-100 (Sigma) and 0.25% BSA (Sigma) (PBS-Triton-BSA), and incubation was conducted at 4 °C overnight. After washing, the sections were incubated with secondary antibodies in PBS-Triton-BSA for 45 min at room temperature. Supplemental Table 1 lists the antibodies used in this study. Slides were counterstained with DAPI (Biotium) for 1 min, and mounted with Dako fluorescent mounting media. Digital images were acquired using the EVOS FL Cell Imaging System (Thermo Fisher), The percent of mice with teratoma was calculated by comparing the number of mice with teratoma to total number of mice transplanted in each replicate (Fig. 2B). The graft area was calculated using ImageJ version https://imagej.net.

### Quantitative PCR

Cells were harvested at day 13 of differentiation and the RNA was extracted using Ambion PureLink RNA mini kit (Invitrogen). RT-PCR was performed to obtain cDNA using Superscript III reverse transcriptase and RNaseOUT recombinant ribonuclease inhibitor (Invitrogen). QPCR was further performed using Bio-Rad SsoAdvanced universal SYBR green supermix on the Biorad CFX Connect real-time PCR system. Relative gene expression was calculated by normalizing mRNA levels of the gene of interested to the housekeeping gene RPL-19, while fold-change was obtained by normalization of the gene expression in cell line compared to the reference sample. The sequences of RT-QPCR primers used are listed in Supplemental Table 2.

## Acknowledgments

This work was supported by funding from the Toronto General and Western Hospital Foundation and from the Canadian Institute of Health Research project grant to M.C.N. Y.A. was supported by post-doctoral fellowships from the JDRF-Canadian clinical trial network, Toronto General Hospital Research Institute as well as Medicine by Design. F.P. was supported by scholarships from the Ontario Graduate Scholarship and the Banting and Best Diabetes Centre. We acknowledge Dr. Amanda Oakie for neurectoderm differentiation, Dr. Stephanie Protze for mesoderm differentiation and Dr. Rangarajan Sambathkumar and Angel Sing for technical support. The graphical abstract and Figures 1A, 1H and 3A were generated using BioRender.com.

## Author contributions

YA designed and conducted experiments, analyzed and interpreted the results and prepared the manuscript. FS designed and conducted experiments and analyzed data. FP assisted with RT-QPCR and *in vivo* GSIS. BN and SNV performed Ku80 staining. ECM generated cells for transplantation. MCN designed the experiments, analyzed and interpreted the results and prepared the manuscript.

