## Supplemental information for "GP2-enriched pancreatic progenitors give rise to functional beta cells *in vivo* and eliminate the risk of teratoma formation"

### Supplemental Fig. 1

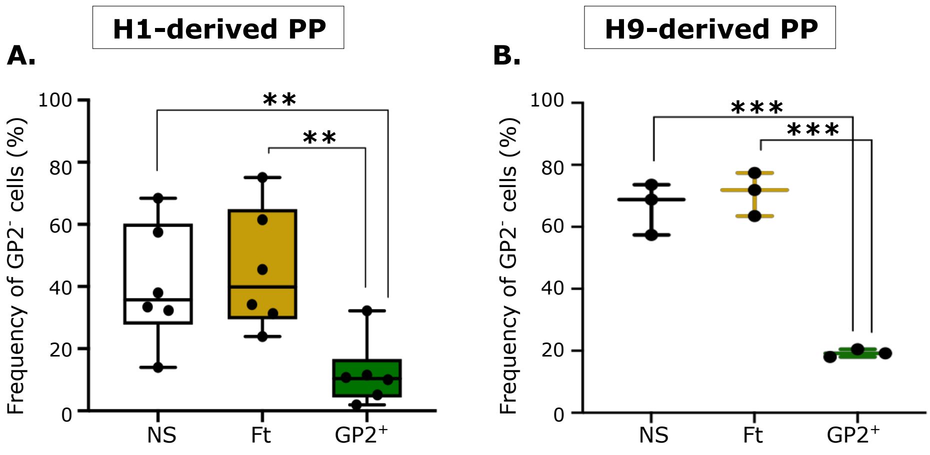


**Supplemental Figure. 1.** Frequency of GP2 negative (GP2$^-$) cells in NS, Ft and GP2$^+$ cell populations after MACS in H1 (A) and H9 (B)-derived cells. (A, n=6; B, n=3; One-way ANOVA analysis with Tukey's multiple test; **P<0.01, ***P<0.0001, error bars represent S.E.M.).

### Supplemental Fig. 2

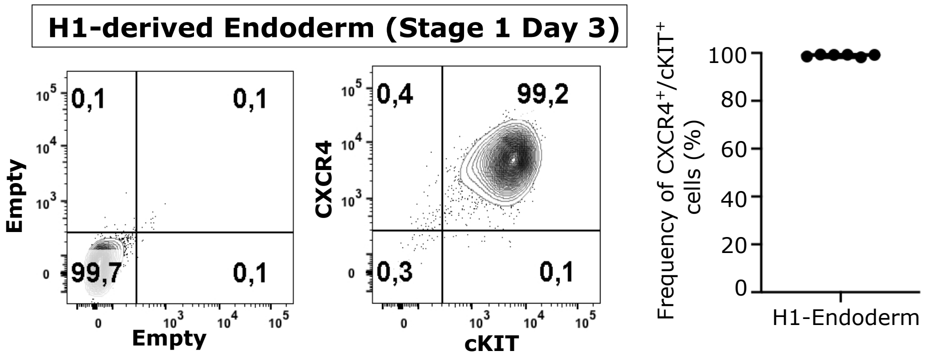


**Supplemental Figure. 2.** Representative flow cytometry plot and quantification of CXCR4 and cKIT in H1-derived endoderm cells at stage 1 (n=6; error bars represent S.E.M.).

### Supplemental Fig. 3

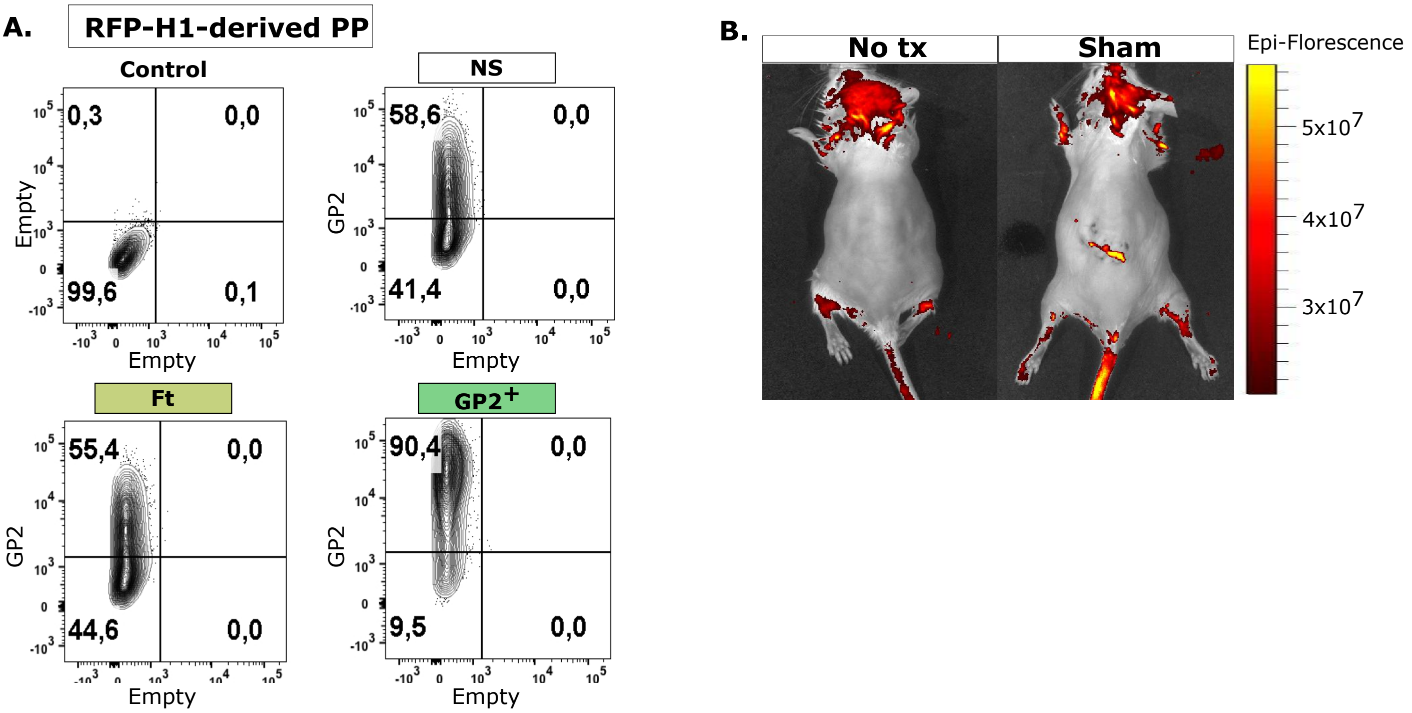


**Supplemental Figure. 3.** (A) Flow cytometric quantification of NXK6-1/PDX1 expression in of RFP-hESC derived PP. (B) Live imaging of mice without transplantation (No Tx), or transplantation of an empty collagen hydrogel (sham) at week 1 post-transplantation.

**Supplemental Figure 4**

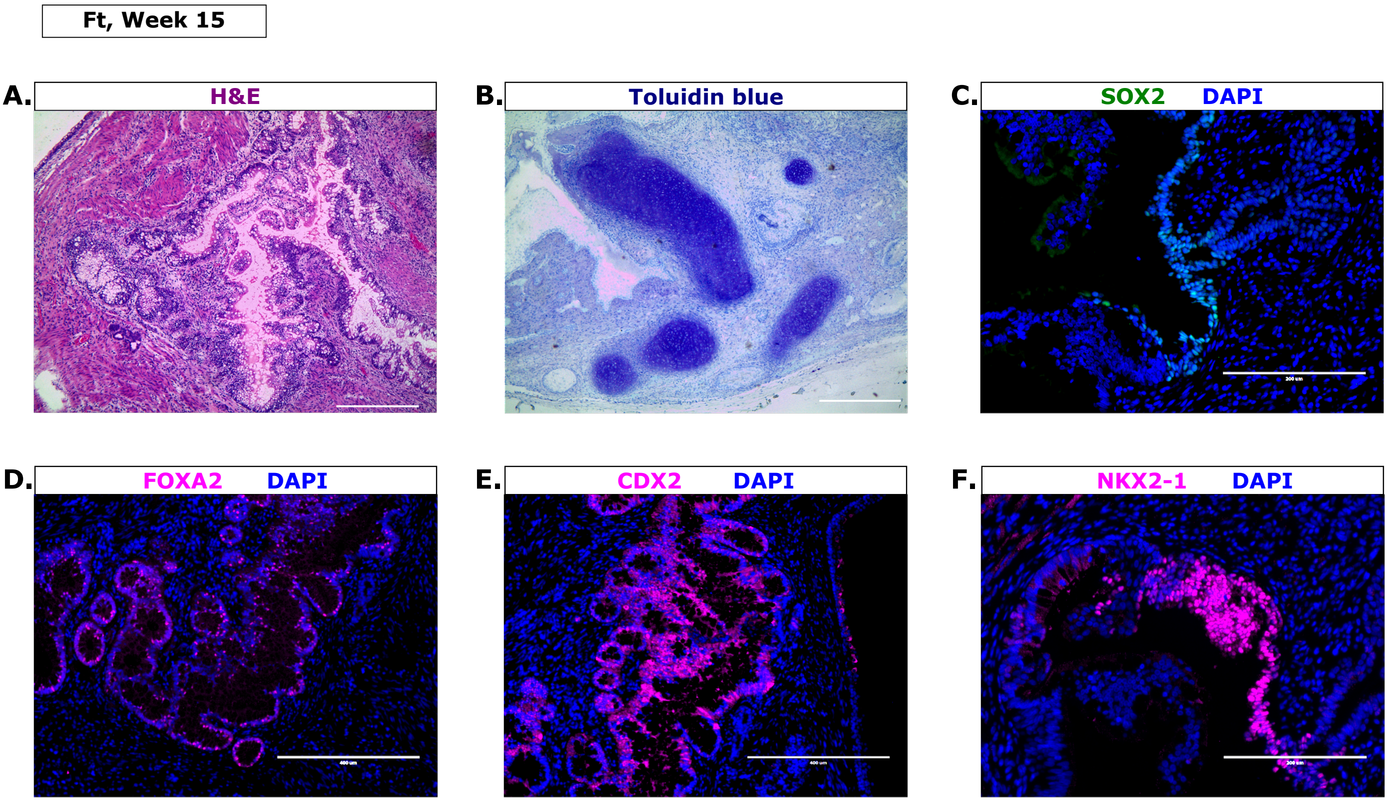


**Supplemental Fig. 4.** (A-F) Representative images of Ft-derived grafts stained with H&E (A), toluidine blue (B), and immunostained using SOX2 (C), FOXA2 (D), CDX2 (E), and NKX2-1 (F) antibodies at week 15 post-transplantation (A, scale bar is 100µm; C, F, scale bar is 200µm; B, D, E, scale bar is 400µm). (J) Percentage of NS, Ft and GP2+-recipient mice that generated teratomas (NS, n=15; Ft, n=17; Ft, n=11).

### Supplemental Tables

**Supplemental Table 1.** List of antibodies used in immunostaining (I) or flowcytometry (F).

| Antigen | Species | Vendor | Dilution |
|---|---|---|---|
| CDX2 | Rabbit | Abcam | 1:300 (I) |
| Chromogranin A (CgA) | Rabbit | Abcam | 1:100 (I) |
| Cytokeratin 19 (CK19) | Rabbit | Abcam | 1:800 (I) |
| C-peptide | Rat | Developmental Studies Hybridoma Bank | 1:1000 (I) |
| FOXA2 | Rabbit | Abcam | 1:500 (I) |
| Glucagon (GCG) | Mouse | Sigma-Aldrich | 1:500 (I) |
| GP2 | Human | MBL | 1:10000 (F) |
| Ki67 | Rabbit | Abcam | 1:1000 (I, F) |
| Ku80 | Rabbit | Cell Signaling | 1:450(I) |
| PDX1 | Goat | Abcam | 1:10000 (I) |
| PDX1 | Goat | R&D Systems | 1:100 (F) |
| Somatostatin (SST) | Rabbit | Abcam | 1:500 (I) |
| SOX2 | | Cell signaling | 1:400 (I) |
| Trypsin | Sheep | R&D Systems | 1:300 (I) |
| NKX2-1 | Rabbit | Abcam | 1: 250 (I) |
| IgG-AF488 | Goat | Jackson Immuno Research Labs | 1:400 (F) |
| IgG-AF488 | Mouse | Jackson Immuno Research Labs | 1:800 (I) |
| IgG-Cy5 | Mouse | Jackson Immuno Research Labs | 1:800 (I) |
| IgG-Cy5 | Rabbit | Jackson Immuno Research Labs | 1:800 (I) |
| IgG-Cy5 | Rat | Jackson Immuno Research Labs | 1:800 (I) |
| IgG-AF647 | Mouse | Jackson Immuno Research Labs | 1:400 (F), 1:800 (I) |
| IgG-AF549 | Mouse | Jackson Immuno Research Labs | 1:800 (I) |
| IgG-AF549 | Rabbit | Jackson Immuno Research Labs | 1:800 (I) |
| R-Phycoerythrin-conjugated AffiniPure Goat Anti-Mouse IgG | Mouse | Jackson Immuno Research Labs | 1:800 (F) |

**Supplemental Table 2.** List of QPCR primers.

| *Gene* | *Forward primer (5' to 3')* | *Reverse primer (3' to 5')* |
|---|---|---|
| *CDX2* | *AGCCAACCTGGACTTCCTGTCATT* | *ACACAGACCAACAACCCAAACAGC* |
| *CELA3* | *ATGACATGCCCCTCATCAAGCTCT* | *ATGTAGCAGGGTGTCTTGTTGGGA* |
| *GP2* | *AACCCTTCCGAAGCACAGAG* | *GGACACAGGTCTCCGACATC* |
| *HNF1b* | *AGA GTA ACA TGC CAG CTT CCT CCT GTG* | *TATCAAACAGCCAGTTTCCCTCCTGCC* |
| *MKI67* | *AATTTGCTTCTGGCTTCCC* | *GAC CCC GCT CCT TTT GAT AGT* |
| *NEUROD1* | *TCC CAT GTC TTC CAC GTT AAG CCT* | *CAT CAA AGG AAG GGC TGG TGC AAT* |
| *NEUROG3* | *GCGCAATCGAATGCACAACCTCAA* | *TTCGAGTCAGCGCCAAGATGTAGTT* |
| *NKX2-1* | *GCCAAACTGCTGGACGTCTTTCTT* | *CCTTGAGATTGGATGCGCTTGGTT* |
| *NKX6-1* | *AGAGGACGACGACTACAATAAGCC* | *ACTTGTGCTTCTTCAACAGCTGCG* |
| *OCT4* | *ATGCATTCAAACTGAGGTGCCTGC* | *CCACCCTTTGTGTTCCCAATTCCT* |
| *PDGFRa* | *GGCAGTACCCCATGTCTGAAG* | *CGTCACAAAAAGGCCGCTG* |
| *PDX1* | *TACTGGATTGGCGTTGTTTGTGGC* | *AGGGAGCCTTCCAATGTGTATGGT* |
| *PTF1A* | *TTATCCGAACAGCCAAAGTCTGGA CC* | *AGTCTGGGACCTCTCAGGACACAA* |
| *RPL19* | *CTCGATGCCGGAAAAACACC* | *TTCTCTGGCATTCGGGCATT* |
| *SOX2* | *GGATAAGTACACGCTGCCCG* | *ATGTGCGCGTAACTGTCCAT* |
| *SOX9* | *TGCATTTCCTCCTGCCTTTGCTTG* | *GGGCACTTATTGGCTGCTGAAACA* |
| *TOP2A* | *GCTGCCCCAAAAGGAACTAA* | *GGCGATTCTTGGTTTTGGCA* |
| *ZIC3* | *TATCAGTCTCGCGCTCAC* | *TGTCTTTTGCGGTTTATCTTCCTG* |
